# Characterization of the KRas G12D-inhibitor interactions by HDX-MS and molecular dynamics simulations

**DOI:** 10.1101/2025.04.22.650091

**Authors:** Evgeniy V. Petrotchenko, Brandon Novy, Edith Nagy, Konstantin I. Popov, Jason B. Cross, Roopa Thapar, Christoph H. Borchers

## Abstract

Hydrogen-deuterium exchange (HDX) combined with mass spectrometry (MS) is a powerful technique for studying changes in protein structure and dynamics upon ligand binding. Protein-ligand complexes can result in increased protection of peptide-bond amides in HDX indicating protein structure stabilization. We have characterized the interaction of lead inhibitor candidates towards the KRas G12D oncoprotein by intact protein and bottom-up HDX-MS in combination with molecular dynamics (MD) simulations. Significant differences in HDX protection were detected upon inhibitor binding in the flexible switch-II pocket of the protein. We identified a correlation between ligand binding affinities and corresponding changes in protection. MD simulations of the free and inhibitor-bound KRas G12D proteins revealed changes in the hydrogen bond network of backbone amides in the switch-II region upon inhibitor binding, explaining the observed HDX protection. This combined HDX-MS and MD analysis approach provides a mechanistic picture of the KRas G12D protein-inhibitor interactions and may be a useful tool for future drug design efforts.

## Introduction

Structural proteomics methods such as limited proteolysis, surface modification, hydrogen-deuterium exchange and crosslinking combined with modern mass spectrometry provide unique experimental data for protein structure determination, characterization of conformational changes, protein-protein and protein-ligand interactions (1). Historically, these approaches have primarily been used for studying proteins and macromolecular complexes, but have great potential for studying protein-ligand interactions as well, making them an attractive methodology for drug development applications. HDX-MS, in particular, is an informative technique for studying protein-small molecule interactions (2, 3). Peptide-bond amide HDX reflects the hydrogen bonding status and the presence of stable secondary structure in proteins. Disordered regions in proteins undergo more efficient exchange and are characterized by higher HDX values. Protein structure can be stabilized by interaction with small molecule ligands, which can manifest as an increase in protection to HDX upon the protein-ligand complex formation. Changes in HDX protection of the target protein upon ligand binding can be measured by intact protein, bottom-up, and top-down MS approaches. Intact protein mass measurements reveal overall stabilization through total deuteration changes, while bottom-up and top-down approaches provide peptide- and residue-specific insights, respectively. These methods identify drug binding sites and allosteric conformational changes by analyzing deuteration patterns upon ligand binding. (4). MD simulations provide insights into protein structure and dynamics and can describe behaviour of the measured by HDX-MS backbone amide hydrogen bonds (H-bonds) (5). Measuring differential HDX between free and ligand-bound protein states can in turn characterize conformational changes upon ligand binding and provide insights into mechanism of ligand binding (6). A combination of such analyses with MD simulations can facilitate modeling of protein-ligand complexes (7) and pinpoint the locations of affected H-bonds, which is critical for the interpretation of HDX-MS data in the drug development and optimization process. Previously, we proposed a simplified empirical approach for determining the opening frequencies of the backbone amide H-bond-forming donor-acceptor pairs along MD trajectories for the interpretation of the mass spectrometry-derived HDX values (8, 9). We expanded this methodology to describe changes in protein H-bond networks upon inhibitor binding and identify structural features in drug-like ligands that drive protein conformational changes.

KRas G12D is one of the most common oncogenic drivers in human cancers. The high prevalence and poor prognosis associated with KRas G12D mutations in various cancers makes it an attractive target for drug design. KRas switch II is a key regulatory region that controls the dynamic transition between the protein’s active and inactive states (10). Targeting this region is a major focus in drug discovery, offering a strategy to develop small-molecule inhibitors that disrupt KRas signaling. In this work, we have characterized the interaction of inhibitors that bind the flexible switch-II pocket of mutant KRas G12D proteins by differential intact protein and bottom-up HDX-MS in combination with MD simulations. HDX-MS analysis was successfully applied for the description of the covalent inhibitors of the KRas G12C (11). Recently, MRTX1133, a potent and highly selective KRas G12D inhibitor was reported (12, 13). Several compounds belonging to the MRTX1133 pyrido[4,3-d]pyrimidine scaffold that bind KRas G12D with a range of affinities have also been disclosed (US Patents WO2021041671, WO2022031678, WO2022132200). Here, we have studied the binding of five compounds from this series to the KRas G12D protein using a combination of HDX-MS and MD simulations. HDX-MS reveals changes in backbone amide H-bonding patterns upon ligand binding, while MD simulations offer a detailed per-residue mechanistic description that complements the HDX data. Additionally, our analysis of the MD trajectories allowed us to extend the characterization of the H-bond network beyond the set identified by HDX. This integrated HDX-MS-MD approach not only enhances our understanding of the protein stabilisation upon ligand binding, but also provides critical structural insights for rational drug design.

## Experimental Procedures

### Protein samples

Recombinant KRas G12D C118S (residues 1-169) protein was expressed in *E*.*coli* BL21 (DE3) RIL cells using a pET28a vector and purified to homogeneity using standard nickel affinity and size exclusion chromatographies. The C118S mutation was previously shown to have no effect on KRas G12D structure, but enhances the stability of Ras proteins (14). 10 mM DMSO stock solutions of compounds were stored until use at -20°C.

### Differential hydrogen-deuterium exchange

For the intact-protein HDX analyses, 2 µL of free or ligand-bound KRas G12D protein samples were mixed with 8 µL of H_2_O or D_2_O (1:4 v:v), incubated for varying time intervals, quenched with 10 µL (1:1 v:v) of 0.2% FA in H_2_O, and 10 µL was immediately injected for LC-MS analysis. LC-MS analysis was performed on a TripleTOF 6600 mass spectrometer (Sciex) interfaced with a Nexera LC system (Shimadzu). Chromatography was performed using short 3-min gradients of 0-100% acetonitrile in 0.1% FA at 200 µL/min flow rate, using an Agilent C_18_ 300Å, 3 *μ*m particle size, 2.1 mm i.d. x 50 mm long column, cooled to 0^°^C in ice bath, MS1 spectra were acquired over the mass range from 100-2000 Da. Analyses were performed in triplicates. Spectra were deconvoluted using PeakView and BioToolKit software (Sciex). Deuteration of the KRas G12D protein was determined based on the mass shift from the mass of the non-deuterated KRas G12D protein. In-exchange was estimated from the deuteration at 0 s, back exchange was estimated from fully deuterated horse myoglobin as ∼30%.

For bottom-up HDX analyses, 2 µL of free or ligand-bound KRas G12D protein samples were mixed with 8 µL of H_2_O or D_2_O (1:4 v:v), incubated for varying time intervals (0, 20, 40, 80 and 160 s), quenched with 10 µL (1:1 v:v) of 0.2% FA in H_2_O, supplemented with 2 µL of freshly prepared 1 mg/mL pepsin solution in H_2_O, placed in an autosampler (for ∼30 s at 10^°^C), and 10 µL was immediately injected for LC-MS analysis. Peptic peptides from the KRas G12D protein were identified from information dependent acquisition (IDA) LC-MS/MS analysis of the protein in water. Data were analyzed by Protein Pilot (Sciex) software. Chromatography was performed as described above. Spectra were analyzed by PeakView software (Sciex). Deuteration of the peptides was determined using MassSpecStudio software package (15). Differential bottom-up HDX protection values were visualized using a blue-white-red palette with red set as the maximum observed Δ% of the corrected deuteration values of the peptic peptides.

### MD simulations and data analysis

The structure of the protein was prepared using Schrödinger Protein preparation tools with default settings (16, 17). The structures of all ligands were prepared using the LigPrep tool to sample all possible conformation states during the docking protocol (18). The best docking models were selected based on the Glide SP docking score, with the top-scored pose chosen as the structural model of the complex (19). MD simulations of the protein-ligand complex models were performed using GROMACS 2020.3 software on UNC high-performance computing clusters with Nvidia V100 GPUs at a constant pressure and temperature of 1 atm and 270 K for 500 ns in three replicates with identical parameters. All replicates were individually minimized and equilibrated to obtain unique initial velocity distributions (20, 21). MD trajectories were processed using the MDTraj package (22) in Python with a timestep of 1 ns for a total of 500 frames. To estimate the changes in the secondary structure, the dynamics of the H-bonds along the trajectory were analyzed by monitoring the distances between backbone H-bond-forming donor-acceptor atom pairs. H-bonds were identified using the Baker-Hubbard algorithm (23), which applies a donor-acceptor distance cutoff of 2.5 Å and a minimum donor-hydrogen-acceptor angle of 120°. The total lifetime of H-bonds were visualized to identify dynamic fluctuations or stability changes, including transient flickering or ligand-induced stabilization. To quantify protection factors and annotate structural features, we aggregated donor occupancy across replicates. H-bonds present in less than 40% of the 500-nanosecond trajectory in either simulation set were filtered out, as these transient interactions would have undergone deuterium exchange and were not relevant for analysis. H-bonds exhibiting significant changes in protection factors were identified using a cutoff of +-20% in occupancy difference between the simulation sets. Principal component analysis (PCA) was performed on the H-bond status of the switch II region to characterize variance in both simulation sets.

## Results and discussion

We used HDX-MS analysis to characterize the binding of the KRas G12D protein with five inhibitors. We observed increased protection from HDX for these compounds. MD simulations of free and ligand-bound proteins corroborated the results, offering an atomic-level view of secondary structure changes in KRas G12D that drive the observed HDX shifts upon ligand binding.

### Changes in HDX protection of KRas G12D protein upon ligand binding

Differential intact and bottom-up HDX-LC-MS analyses were performed to identify conformational changes of the KRas G12D protein upon inhibitor binding. For differential HDX analysis, free and ligand-bound protein samples were exposed to D_2_O solution for varying time intervals, and the deuteration levels were measured by mass spectrometry. Differences in total deuteration between free and ligand-bound KRas G12D reflect the overall stabilization of its secondary structure upon complex formation. Analysis of the differences in deuteration of the peptides provides the locations of differentially protected regions upon complex formation and therefore can suggest the location of the inhibitor-binding site.

Intact HDX analysis of free and inhibitor-bound Kras G12D samples revealed a significant increase in protection of ∼11 protons upon formation of the complexes, at the 20 s exchange time (Figure 1). Previously, we showed that this exchange time has a good correlation between measured deuteration values and number of the H-bonds in crystal structures of folded proteins (24). Observed values can serve as a good estimation of number of H-bond-forming donor-acceptor atom pairs undergoing changes upon protein-ligand complex formation. Such analysis is quite precise (%CV<3), fast and can be adapted as a screening method for lead drug-like candidates. We observed some subtle differences in differential exchange for compounds of differing affinities (Figure 1D).

**Figure 1.**
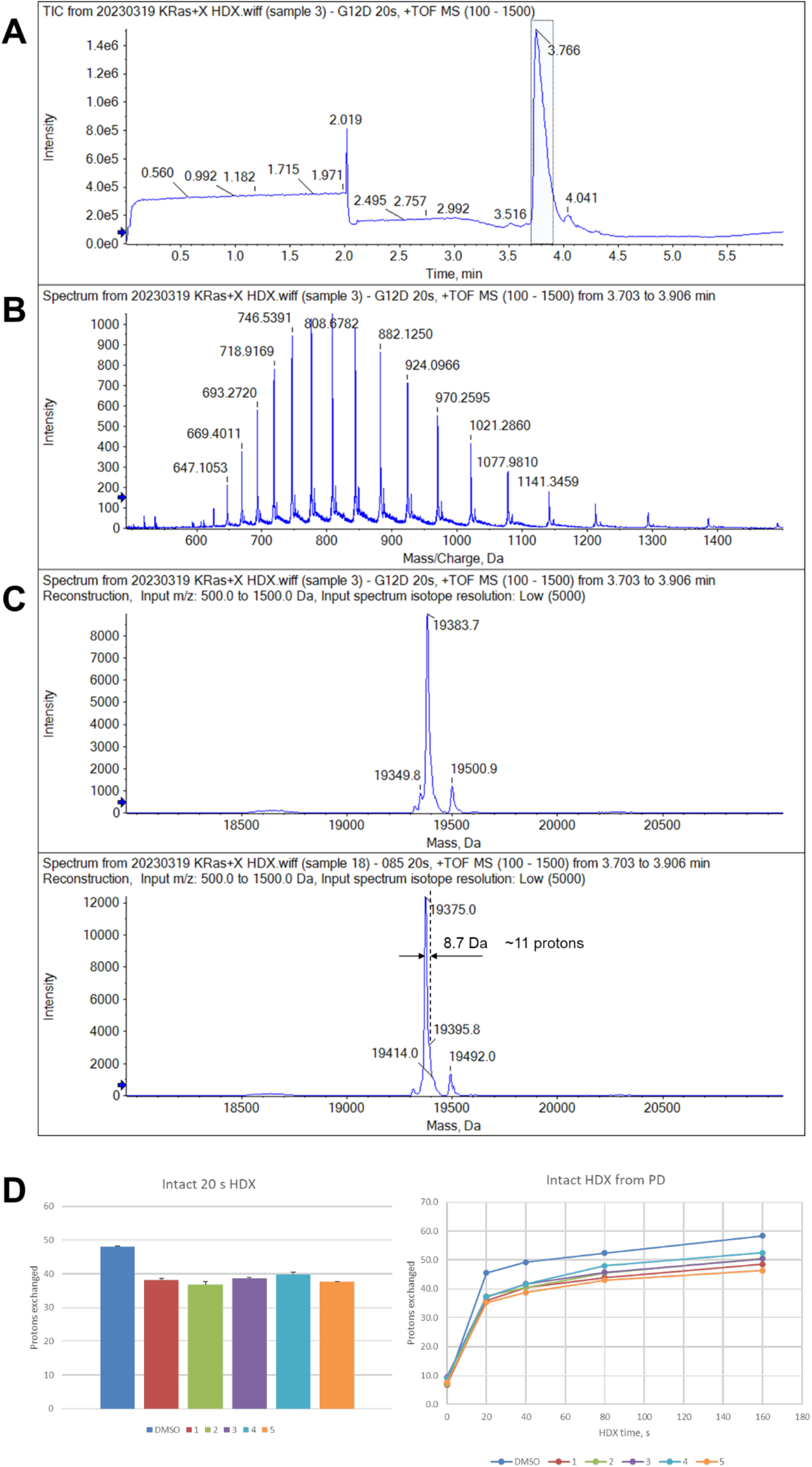
Intact protein mass HDX-LC-MS analysis of the KRas G12D protein – drug interactions. **A**. Representative total ion chromatogram. **B**. Representative cumulative mass spectrum. **C**. Deconvoluted cumulative mass spectrum of free (top) and ligand-bound (bottom) KRas G12D protein. **D**. Changes in KRas G12 HDX protection for different compounds. Left, intact mass 20-s HDX (n=3); right, kinetic plots from intact mass HDX.

Differential bottom-up HDX analysis of free and ligand-bound KRas G12D protein samples also revealed a few peptides exhibiting increased protection against exchange upon ligand-KRas G12D complex formation (Figure 2). Visualization of the observed changes on the three-dimensional structures of the KRas G12D allowed us to locate the regions which showed a change in protection upon inhibitor binding. The regions that exhibited the highest changes involved loops which are flanked by secondary-structure elements. The changes in protection were ligand specific. Notably, these differences were similar for both bottom-up and intact HDX analyses. The extent of changes in protection upon complex formation correlated with the inhibitor binding affinities (25). The observed changes were not due to differences in occupancies, as all inhibitors used in the HDX experiment saturated the protein by over ∼97% under the given conditions (Figure S1). Changes in HDX upon ligand binding were mainly detected in the peptides spanning the switch II region. This is consistent with the location of the inhibitor binding site in the cleft between the switch II loop and the core of the protein, as was observed in the crystal structure of the KRas G12D – Compound 5 complex. The observed changes in HDX protection upon protein-inhibitor complex formation would then suggest an increase of secondary structure content in the switch II loop.

**Figure 2.**
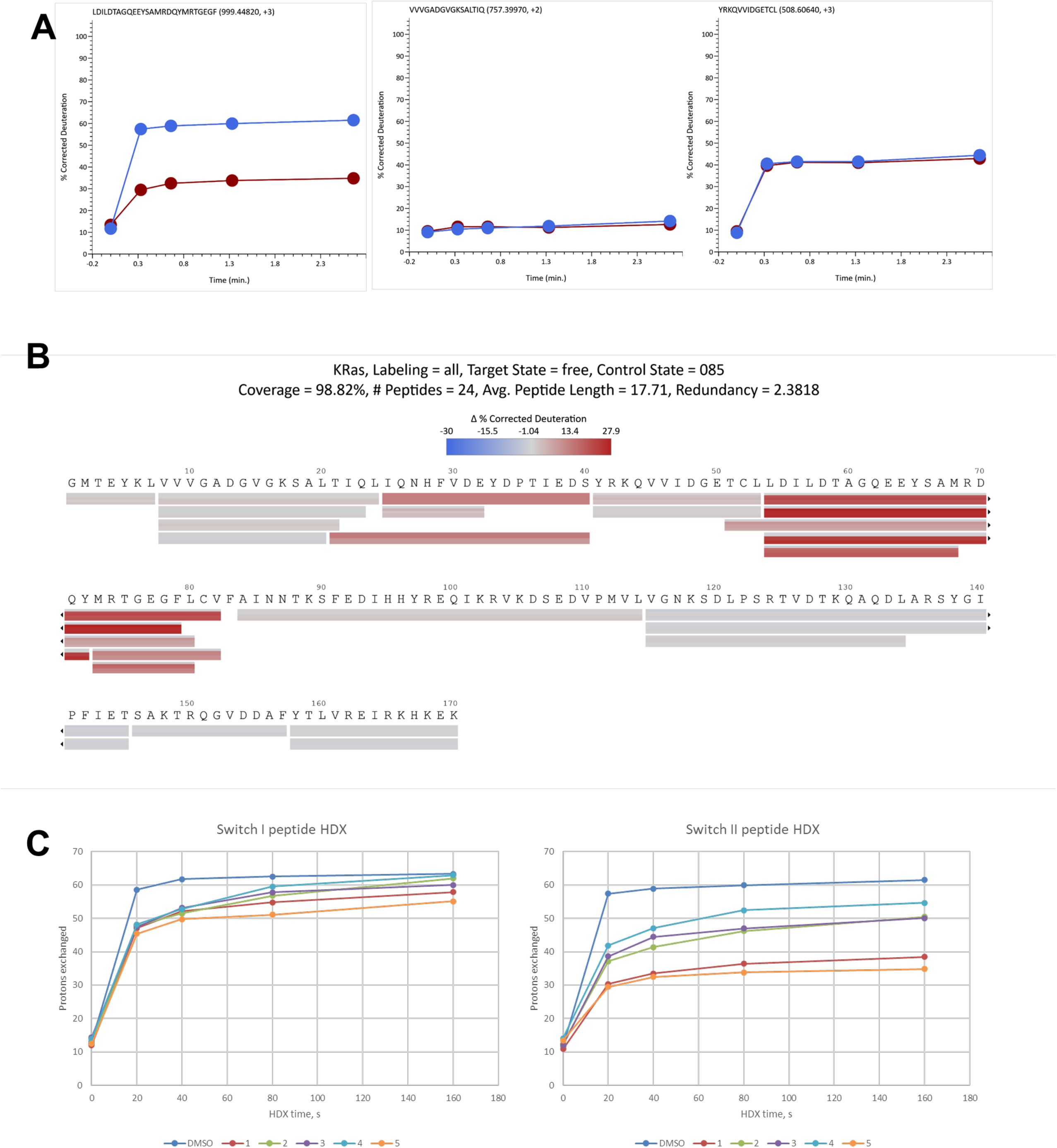
Bottom-up HDX-LC-MS analysis of the KRas G12 protein – drug interactions. **A**. Representative kinetic plots of KRas G12 peptic peptides HDX from left to right: differentially protected upon the complex formation peptide, similarly protected slow-exchanging and fast-exchanging peptide. **B**. Differences in deuteration of the peptic KRas G12D peptides between free and Compound 5 drug-bound KRas G12D samples. Values are presented in blue-white-red palette, with red representing higher protection values in KRas G12D-ligand sample compared to the free KRas G12D sample. Bars represent peptides. Slices of bars from top to bottom correspond to 0, 20, 40, 80 and 160 s HDX time points. **C**. Changes in KRas G12D HDX protection for different compounds. Left, kinetic plots of switch I peptide; right, kinetic plots of switch II peptide.

### Molecular dynamics interpretation of the experimentally observed HDX values

Measured HDX values correspond to the exchange of the proton from the nitrogen atom of the backbone amide with deuteron from the solvent. Molecular dynamics simulations in an aqueous environment allow tracking the behavior of each hydrogen atom in the protein throughout the trajectory. A common simplified assumption is that H-bonded backbone amide protons either do not exchange with solvent protons or do so at a significantly slower rate than non-bonded hydrogen atoms. The extent of this exchange will depend on the fraction of time a given amide hydrogen is in a bonded versus free state. By monitoring the states of selected peptide bond amide hydrogens along the simulation trajectory, we can quantify the number of frames at which they exist in a free or bonded state. Hydrogens that spend more frames in the free state correspond to highly exchangeable protons, whereas those that remain bonded for a greater number of frames correspond to protected hydrogens (5). Previously, we showed good correlation between frequencies of H-bond opening for peptide bond amides with observed HDX values (8). Here, we have extended this semi-empirical metric for the characterization of experimentally determined HDX values. Total number of the backbone amides potentially forming H-bonds was estimated from the crystal structure of the KRas G12D - MRTX1133 complex (PDB ID: 7RPZ) as 102 (∼60% of total 170). In free KRas G12D, 69 amides were estimated to be deuterated at 20 s exchange time, which corresponds rather well to the number of non-bonded (based on crystal structure) backbone amide protons. Ligand binding results in an increase of protection for ∼11 protons at 20 s exchange time (Figure 1).

Molecular dynamics trajectory analysis of both the free and ligand-bound states revealed that several H-bonds in the switch II region of KRAS G12D are stabilized in the presence of Compound 5 (Figure 3, S2A). To analyze and compare H-bond networks between liganded and unliganded simulations, considering the network configurations in every frame along each trajectory, we applied principal component analysis (PCA) for dimensionality reduction. The variance of the first two principal components reflects the dynamics of H-bond formation and breaking within the network. In the liganded case, the variance is smaller, indicating more structured and stable hydrogen bond dynamics along the MD trajectories. In contrast, the unliganded simulations show greater variance, suggesting more dynamic hydrogen bond networks, which implies higher conformational flexibility of the unbound protein. (Figure S2B). For the entire protein, we identified a total of 91 H-bonds by aggregating acceptors into a single H-bond (Figure 4A), which agrees well with H-bond counts in HDX-MS data and crystal structure estimates. All identified H-bonds were mapped on a per-residue basis onto the protein structure and displayed in Figure 4B. Eight H-bonds displayed substantial shifts in protection (Figure 4A) across our replicate simulations. Notably, our analysis confirms that most of the significant protection changes are located within the switch II region where ligand binding increases protection. These results align well with the experimental HDX-MS data. For example, the Asp69-Ser65 H-bond forms upon residue coordination with the ligand’s hydroxyl group stabilizing the switch II loop. In the absence of the ligand, this region becomes more dynamic, exhibiting reduced H-bond coordination. (Figure 3, Figure 5). In general, ligand binding stabilizes the switch II residues and restricts loop movement, increasing the chance that H-bonds within the loop will form and remain intact. This leads to a higher degree of protection from proton exchange, as measured by HDX-MS. (Figure 5).

**Figure 3.**
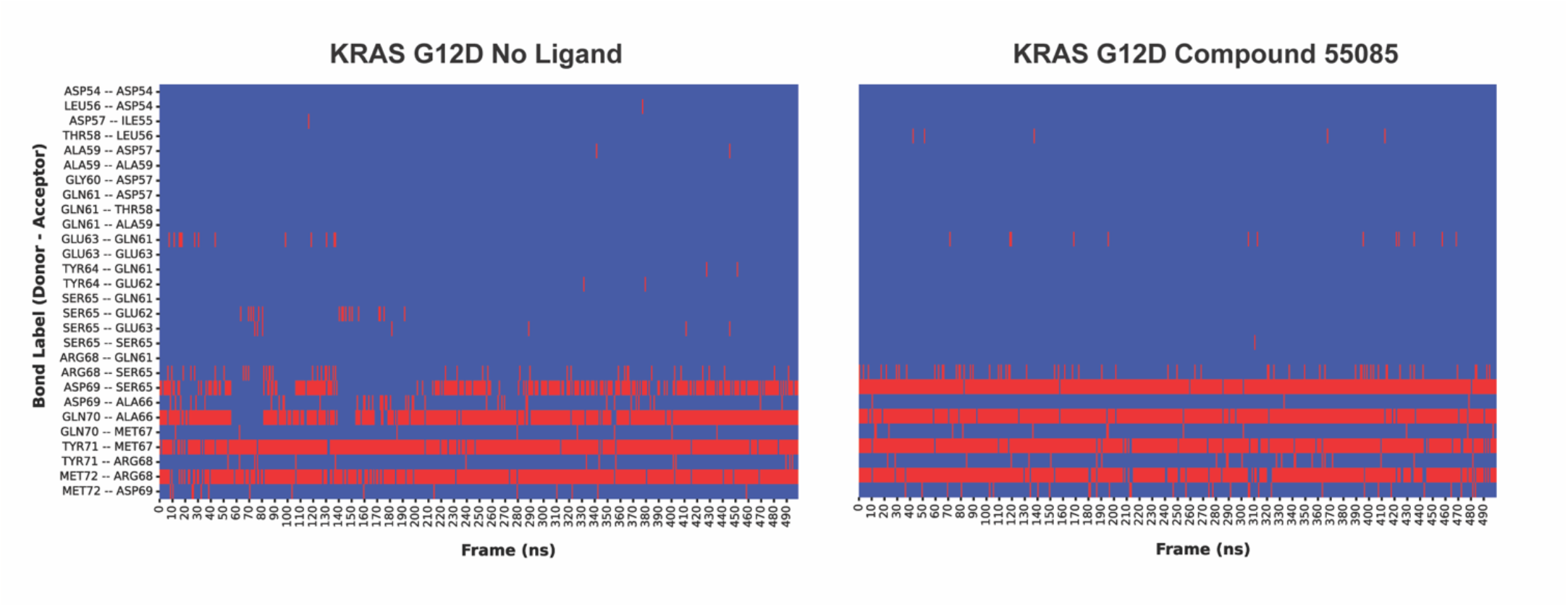
Changes in backbone amide hydrogen bonding at the switch II region observed in MD trajectories upon KRas G12 protein-ligand binding. Transient hydrogen bonds in ligand-free state (left) are stabilized upon ligand binding (right), as illustrated by increased number of frames with occupied H-bonds (shown in red).

**Figure 4.**
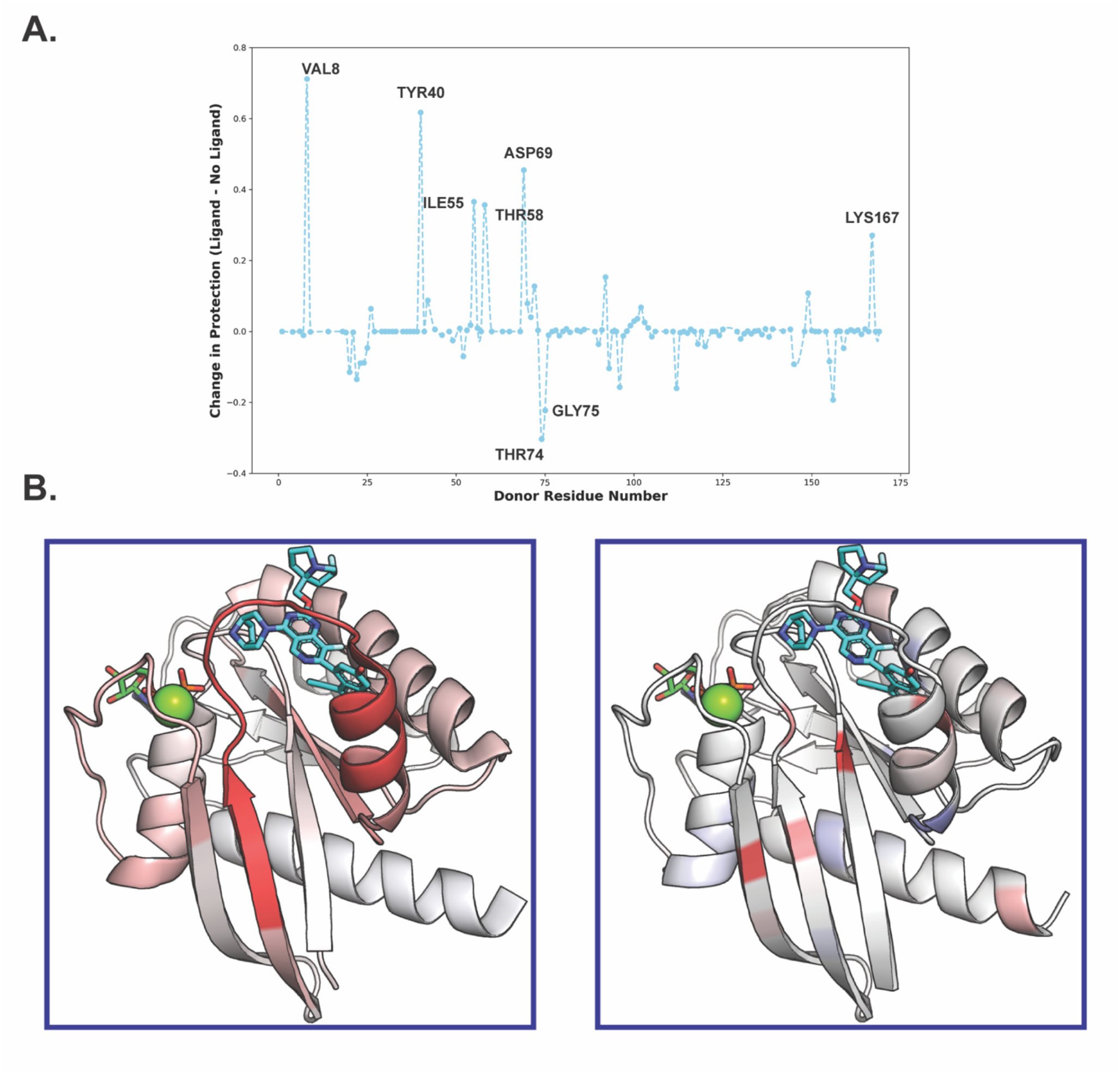
Changes in HDX protection upon KRas G12 protein-drug binding from HDX-MS and MD approaches. **A**.H-bonds above zero show increased protection upon ligand binding. To include H-bonds observed in simulations but below the lifetime threshold, their donor residues were imputed to zero (91 H-bonds above cutoff in either set, 131 total donors displayed). **B**. Visualization of deuteration differences (left) and H-bond occupancy from MD simulations (right) between free and Compound 5-bound KRas G12D. The values are plotted on the KRas G12D-Compound 5 3D structure (PDB: 7RPZ) using a blue-white-red palette, with red indicating higher protection values and stable H-bonds in the KRas G12D-ligand complex compared to the free KRas G12D.

**Figure 5.**
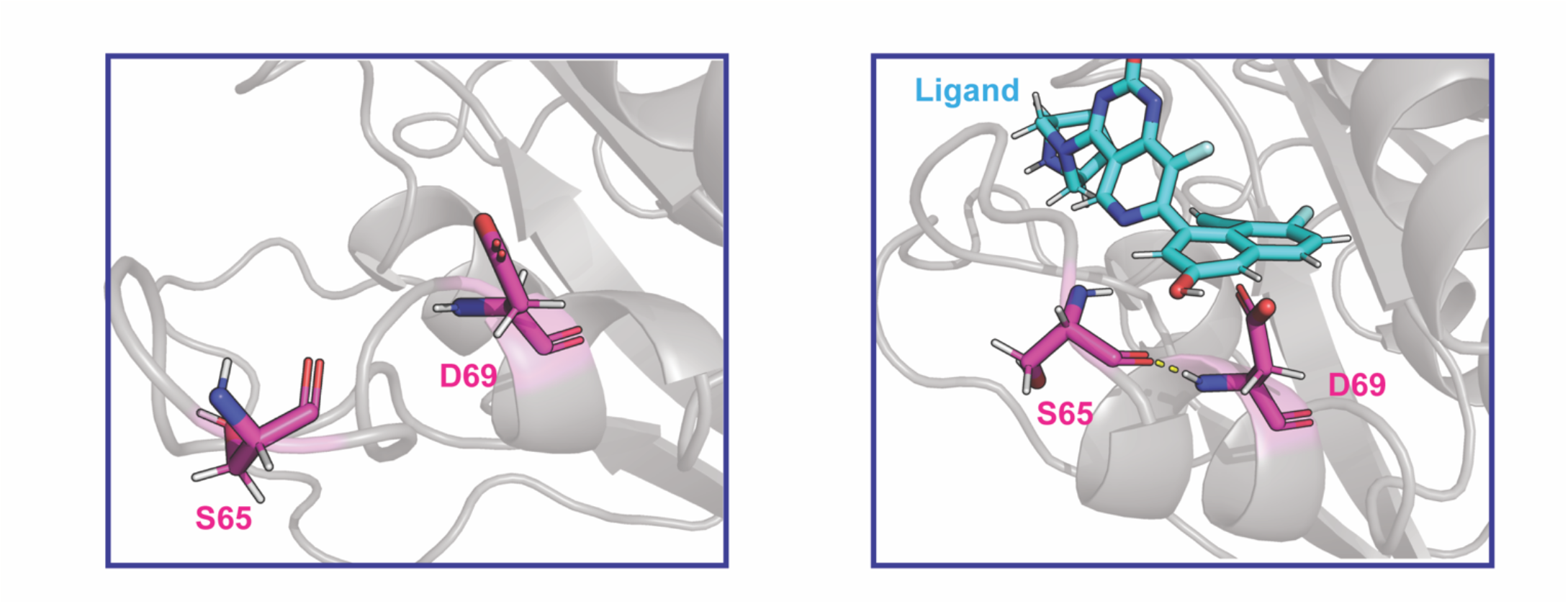
Representative example of hydrogen bond formation within switch II region upon KRas G12-ligand binding. Left, ligand-free state; right, ligand-bound state.

Overall, the combined HDX-MS and MD analyses explain the experimentally observed HDX protection changes upon inhibitor binding and provide atomic-level insights into the mechanisms of protein conformational changes.

## Conclusions

We have characterized ligand-induced stabilization of the protein secondary structure by HDX-MS-MD. We propose semi-empirical metrics for observed experimental HDX values based on H-bonding status from MD trajectories. The analysis provides atomistic details of the H-bond network and interpretation of the experimental data and can be generally applicable for other protein-ligand systems.

## Abbreviations

HDX-MS: hydrogen-deuterium exchange combined with mass spectrometry
MD: molecular dynamics
LiP: limited proteolysis.

## Acknowledgments

This work was supported by funding to CHB for “The Metabolomics Innovation Centre” from Genome Canada through the Genomics Technology Platform (265MET and MC4T), and by grant #42495 to CHB from the Canadian Foundation for Innovation via the Major Sciences Initiatives Fund. CHB is also grateful for support from the Segal McGill Chair in Molecular Oncology at McGill University, and for support for the Segal Cancer Proteomics Centre at the Jewish General Hospital (Montreal, Quebec, Canada) from the Warren Y. Soper Charitable Trust and the Alvin Segal Family Foundation. The authors gratefully acknowledge support from the NIH Biophysics Training Grant (T32GM148376-01A1).

