## Supplemental Material for "Characterization of the KRas G12D-inhibitor interactions by HDX-MS and molecular dynamics simulations"

**Supplementary data**

Figure S1. Compounds used in the study.

Figure S2. Hydrogen bonding status of the backbone amides of the KRas G12D switch II region in free and ligand-bound states.

Figure S3. Hydrogen bonding status of the backbone amides of the KRas G12D protein in free and ligand-bound states.

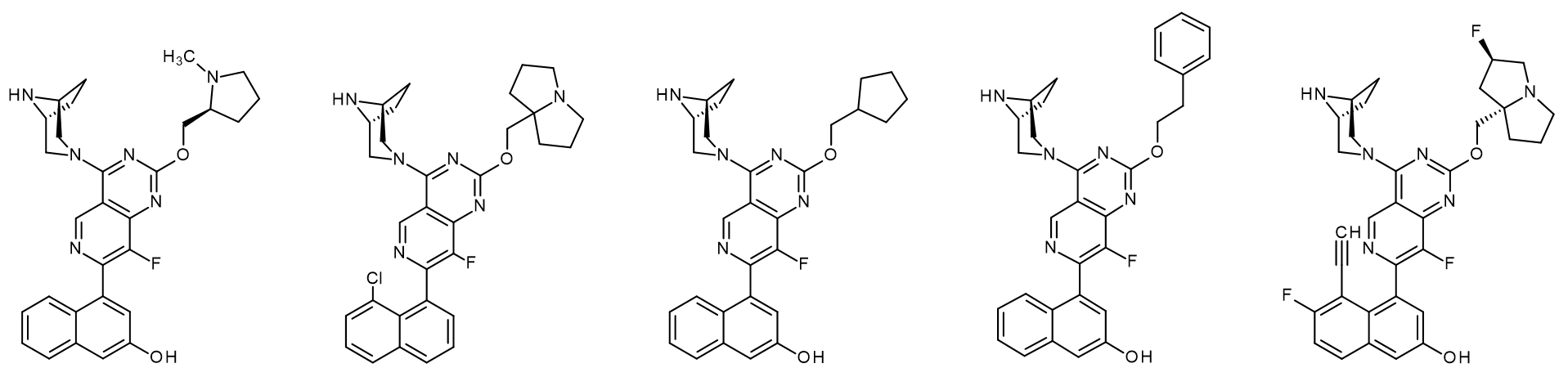
**A**

1	2	3	4	5

**B**

| Compound | P0, nM | L0, nM | Kd, nM | PL, nM | Pf, nM | Pf, % |
| --- | --- | --- | --- | --- | --- | --- |
| 1 | **2000** | **20000** | **100** | 1988.96 | 11.04 | 0.55 |
| 2 | **2000** | **20000** | **5.5** | 1999.39 | 0.61 | 0.03 |
| 3 | **2000** | **20000** | **573** | 1938.50 | 61.50 | 3.07 |
| 4 | **2000** | **20000** | **710** | 1924.41 | 75.59 | 3.78 |
| 5 | **2000** | **20000** | **0.0002** | 2000.00 | 0.00 | 0.00 |

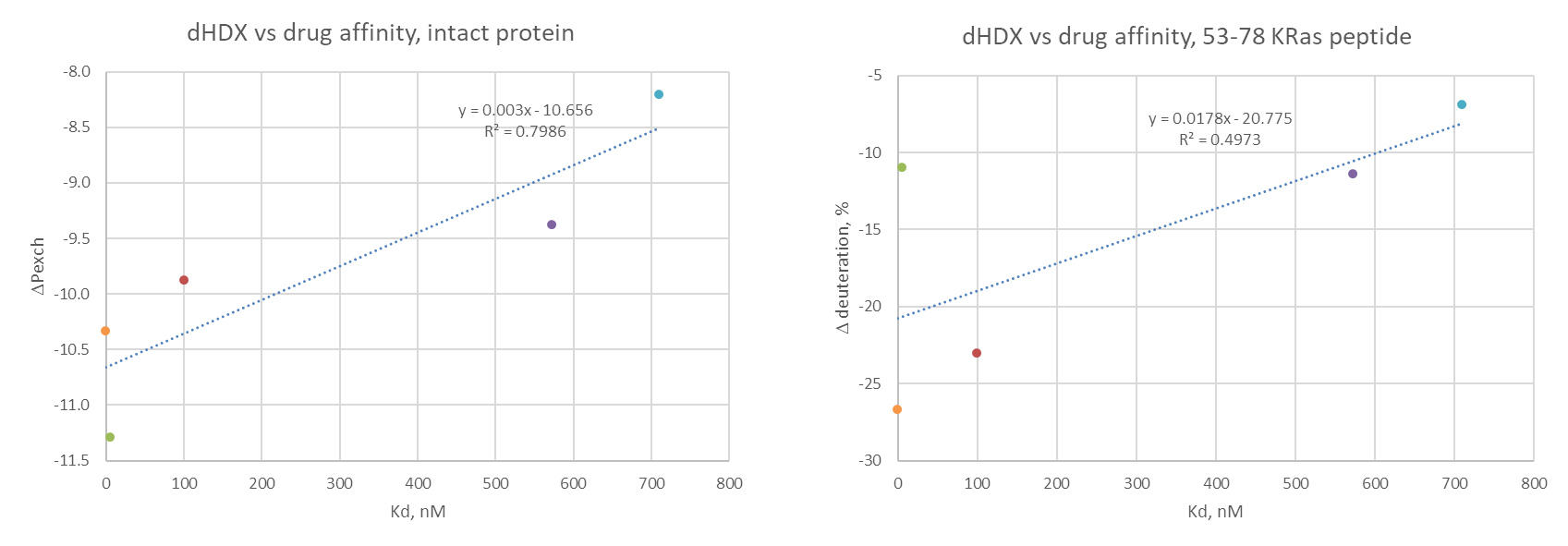
**C**

**Figure S1. Compounds used in the study. A.** Chemical structure of the compounds used in this study. **B.** Theoretical concentration of the free and bound forms of the KRas G12D protein in the HDX reaction mixtures. **C.** Correlation of the observed changes in HDX protection upon protein-ligand complex formation with binding affinities of the compounds.

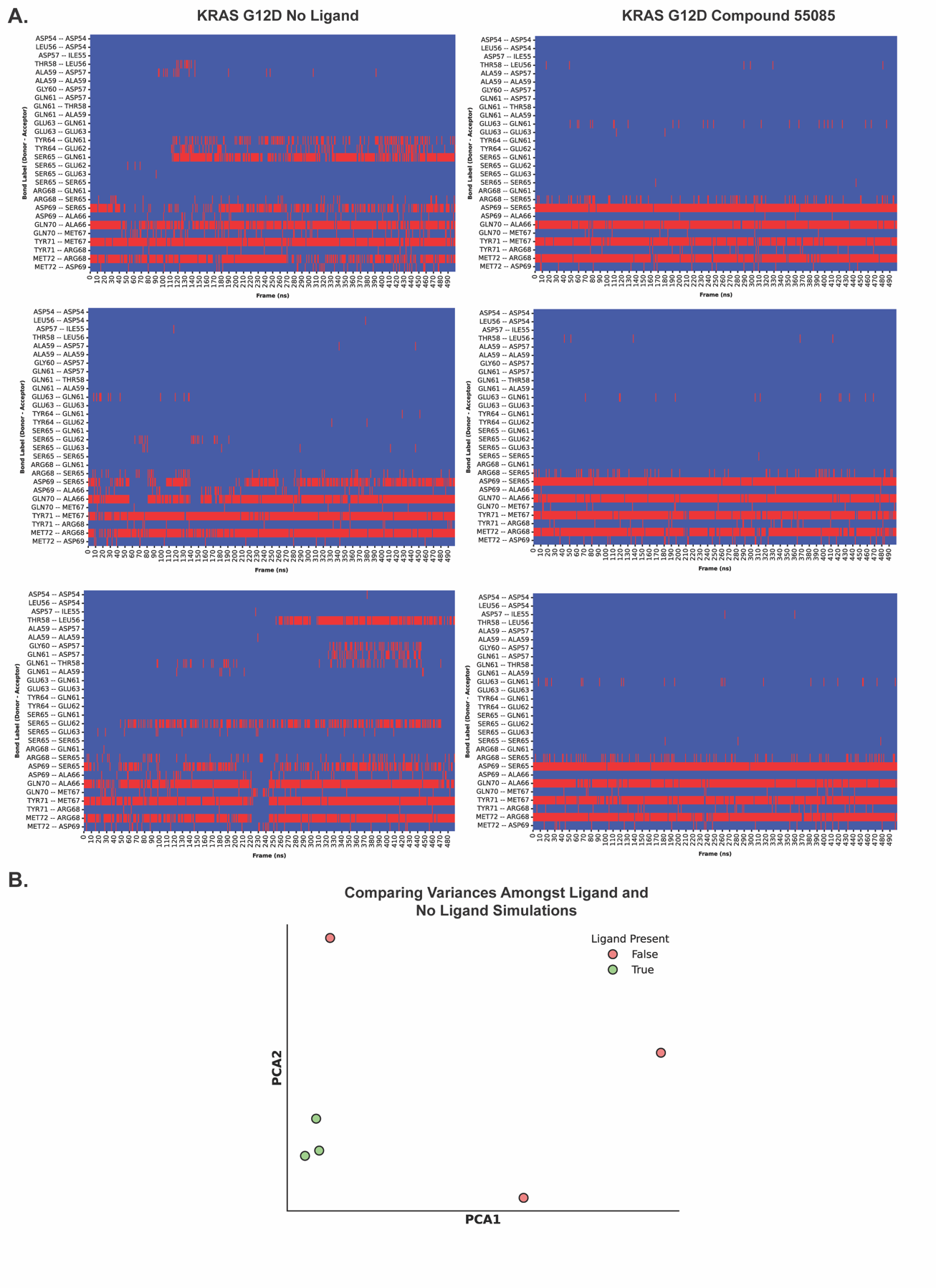

**Figure S2. Hydrogen bonding status of the backbone amides of the KRas G12D switch II region in free and ligand-bound states. A.** H-bonds from MD simulation trajectories for free (left panel) and Compound 5-bound (right panel) states in switch II region (50 -70) are shown. **B.** Principal component analysis (PCA) for H-bond network comparison between liganded and unliganded simulations. The variance of the first two principal components reflects the dynamics of H-bond formation and breaking within the network.

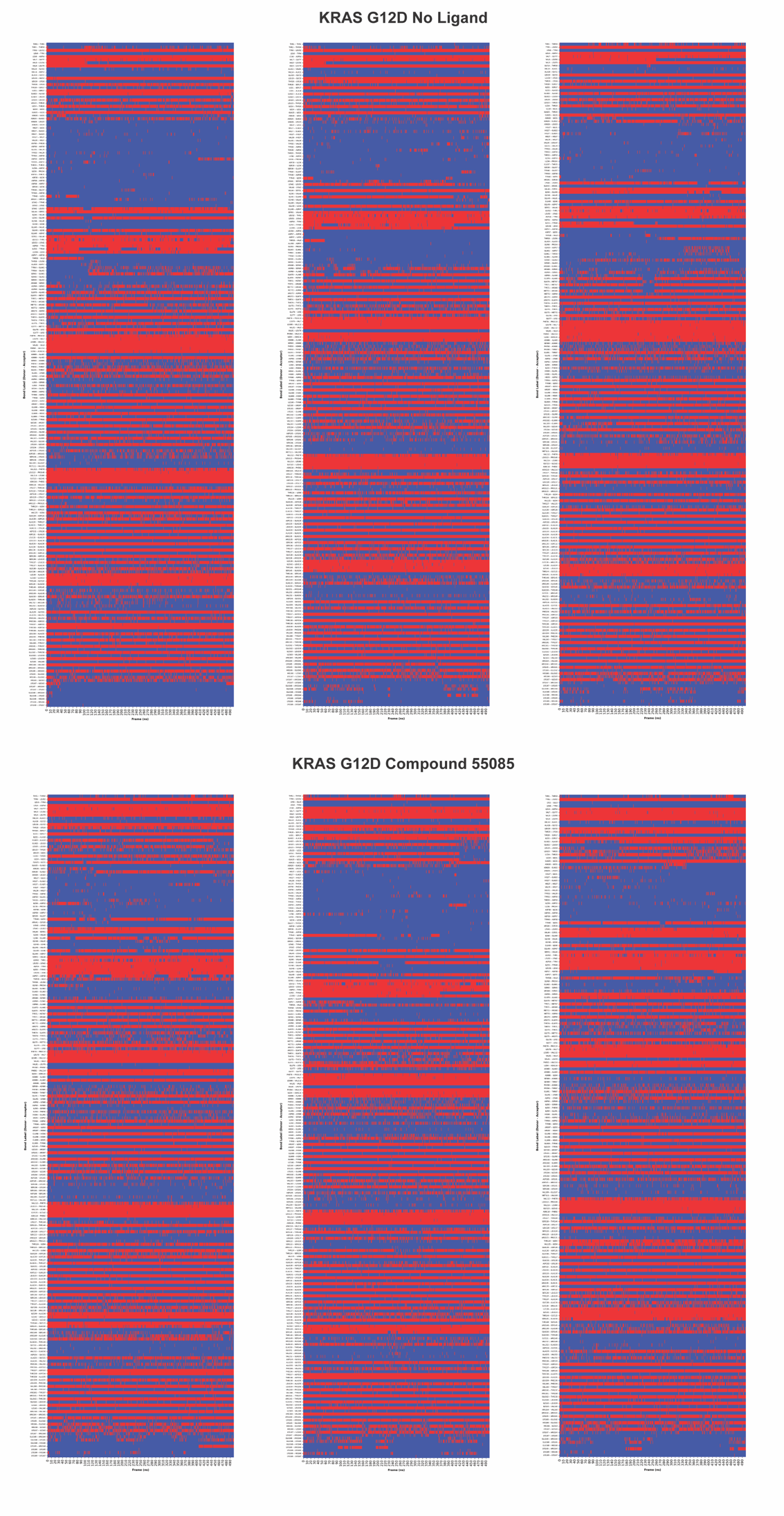

**Figure S3. Hydrogen bonding status of the backbone amides of the KRas G12D protein in free and ligand-bound states.** H-bond occupancies from MD simulation trajectories for free (top panel) and Compound 5-bound (bottom panel) states are shown.
